# Imaging [^18^F]FDG PET/CT Study of Nicotinic Acetylcholinergic Receptor α2 Knock-Out Mice and α2 Hypersensitive Mice Compared to Control Mice: Male-Female Differences and Nicotine Effects

**DOI:** 10.64898/2026.03.23.713331

**Authors:** Christopher Liang, Titus E. Tucker, Ashlee D.L. Coronel, Ellen H.N. Nguyen, Janet L. Nguyen, Irakli Intskirveli, Ronit Lazar, Raju Metherate, Jogeshwar Mukherjee

## Abstract

**Objective:** Nicotinic acetylcholinergic receptors (nAChRs), comprising of α and β subunits are present in the brain and whole body. The less abundant α2-subunit is a fast-acting receptor subtype and plays an important role in cognition and learning. To understand cellular functional consequences, this study evaluated glucose metabolism using [^18^F]FDG PET/CT in α2 knockout (α2KO) and α2 hypersensitive (α2HS) mice.

**Methods:** Control (CN; 4M, 4F), α2 knockout (α2KO; 4M, 4F) and α2 hypersensitive (α2HS; 4M,4F), 12-16 month old mice were used. Mice were fasted and injected with [^18^F]FDG (3-5 MBq) while awake. After 40 minutes they underwent whole body PET/CT. On a separate day, nicotine challenge [^18^F]FDG studies were done. Reconstructed images were analyzed to obtain standard uptake values (SUV) of [^18^F]FDG in brain and interscapular brown adipose tissue (IBAT). Statistical analysis was performed.

**Results:** The α2HS male mice exhibited the largest brain increase in [^18^F]FDG compared to α2KO male mice. The rank order of brain [^18^F]FDG uptake in the 3 groups: α2HS♂> CN♂> α2KO♂> CN♀= α2KO♀≥ α2HS♀. Nicotine treatment reduced brain [^18^F]FDG uptake in all mice. Females had lower [^18^F]FDG uptake compared to males and were less sensitive to α2 nAChR. In the case of IBAT, α2KO mice had significantly higher baseline [^18^F]FDG uptake compared to the other two groups: α2KO♂> α2KO♀> α2HS♀> α2HS♂> CN♀> CN♂. Nicotine decreased IBAT in α2KO mice rather than increase as observed in CN and α2HS mice.

**Conclusions:** α2 nAChRs plays a significant role in brain activation as exhibited by the increase in [^18^F]FDG in α2HS mice. In the absence of regulatory control by the α2 nAChR, the α2KO mice IBAT exhibited higher [^18^F]FDG IBAT compared to controls and α2HS mice. Female mice were less affected by nicotine compared to the male mice. Overall, α2 nAChRs played a significant role in glucose metabolism in the brain and IBAT.

## 1. Introduction

The heteromeric α4β2* nAChRs (the asterisk denotes the various subtypes including α2, α3 and α4 subunits) have a high affinity for nicotine and mediate behaviors related to nicotine addiction and thus serve as targets for nicotine cessation agents [1, 2]. Other subunits that have been implicated in nicotine-related disorders include the α2, α3, α5 and β4 nAChR subunits [3, 4]. The α2 nAChRs have attracted much attention because of their potential role in cortical function [5]. Studies with monkeys showed that the pattern of high-affinity nicotine binding better matched α2 expression than α4 expression [6]. Higher affinity of nicotine and the positron emission tomography imaging probe, [^18^F]nifene exhibited higher affinity for α2 compared to α4 nAChRs [7]. Prior studies of the α2 subunit have also shown that it plays a role in hippocampus-dependent learning and memory and potentiates several nicotine-modulated behaviors [8]. Mice with a hypersensitivity mutation of the α2 nAChR showed a deficit in fear learning which was rescued with low-dose nicotine. On the other hand, mice without the α2 nAChR exhibited learning deficits that were not rescued by nicotine administration [9]. Although the α4 nAChR has been studied more extensively than the α2 subunit, it has been shown that α2β2 nAChRs have similar responses to nicotine, positive allosteric modulators, and partial agonists of the nAChR as do α4β2 nAChRs [10]. Therefore, it appears worthwhile to study the α2 nAChR more in depth and elucidate further its role in disease models [11, 12] and nicotine modulated behaviors [13].

The habit-forming properties of nicotine acting via brain nAChR subtypes are thought to be related not only to its reinforcing properties, but also to its effects on appetite, attention, and mood [14, 15]. Acute nicotine administration has been associated with improved short and long-term memory in mice [4], whereas chronic or prenatal exposure has been shown to provoke behavior that suggests decreased attention [<u>16</u>, <u>17</u>). Altogether, the literature suggests that the different brain nAChR subtypes are important in a variety of cognitive pathways related to nicotine.

Nicotine has been shown to affect cerebral metabolic rates in human brains (measured using [^18^F]FDG PET), suggesting resemblance with other drugs of abuse in reducing overall brain metabolism [18, 19]. It has been suggested that nicotine enhances neurotransmission through cortico-basal ganglia-thalamic circuits directly (via nAChRs) and indirectly (via dopaminergic system) and other mechanisms [20]. This may cause differential effects of glucose metabolism in different brain regions, with increases in some brain regions while decreasing in others. Electrophysiology findings suggest that in the presence of nicotine, α2 subtype may exhibit sustained activation and may play a key role in regulating cortical function [5]. Effects of nicotine on glucose metabolism via the α2 nAChR subtype remains unknown.

Glucose metabolism has also been affected by nicotine in brown adipose tissue (BAT) [21, 22]. BAT functions via thermogenesis by combusting lipids and glucose when active [23] and induced by β3 adrenergic drugs [24, 25]. Studies have shown that combinations of nAChR α3 with β2 or β4 subunits are most common in the sympathetic nervous system which controls adipose tissue activity [26]. Implicated nAChR subunits that have been found within BAT itself are α2>β2>β4, while in white adipose tissue (WAT) the expression is α2>α5>β2 [27]. The β2 subunit has been shown to play a role in nicotine-mediated expression of specific adipokines while the α2 subunit has been found to have importance in the function of beige-like adipocytes [28]. Specifically, activation of beige-like adipocytes results in induction of the α2 nAChRs in subcutaneous fat and inactivation of this receptor led to increased obesity [28, 29].

In an effort to understand the possible effects of the α2 nAChRs on glucose metabolism in the brain and BAT, we have studied three groups of mice using [^18^F]FDG PET/CT. These include C57BL/6 wild type (CN), α2 knock-out (α2KO) and α2 hypersensitive (α2HS) with both male and female mice. Baseline studies (without nicotine) and nicotine pretreatment imaging studies were carried out and compared across the three groups.

## 2. Materials and Methods

### 2.1 Mice

This study was conducted under protocols approved by the University of California Irvine Institutional Animal Care and Use Committee protocol # AUP-25-068, approval period 6/13/2025 to 6/13/2028.

Table-1 summarizes the mice used in this study. Control mice (CN) were C57BL/6 strains. For α2 mice, two mouse lines with α2 nAChRs deleted (*Chrna2* -/-) for α2KO or made hypersensitive (α2HS) with a single substitution of a serine for a leucine (L9’S) (*Chrna2* L9’S/L9’S) were used. The *Chrna2* L9’S/L9’S mutation produces a 100-fold increase in sensitivity to acetylcholine in oocytes [8]. The reported EC_50_ values in the presence of acetylcholine were 3.7 μM for α4β2 nAChR and for the mutant α2β2 nAChR EC50 was 0.041 μM [8]. In rat brain slices, acetylcholine had an affinity of Ki= 19.2 nM for α4β2* nAChR’s in the presence of physostigmine (inhibitor of acetylcholinesterase) and without physostigmine the affinity was Ki= 34.7 μM [30]. Selective subtype affinity of nicotine for α2β2 Ki= 3.62 nM, α3β2 Ki= 8.46 nM and α4β2 Ki= 5.45 nM nAChR in human cloned receptors [7]. The α2HS mice would therefore be a good model to enhance functional activity mediated via the α2β2 nAChR because of this selectivity over other subtypes for acetylcholine and nicotine.

**Table-1:** Summary of mice used in the study.

| Mice Type | Age, month | Weight | [<sup>18</sup>F]FDG MBq, ip | Brain [<sup>18</sup>F]FDG Scans<br>BAT [<sup>18</sup>F]FDG Scans | BAT/WAT CT |
| --- | --- | --- | --- | --- | --- |
| **Control Mice (CN)** |  |  |  |  |  |
| C57BL (WT) Male<br>n=4 | 12 mo | 36 - 46g<br>34 - 36g | 4.22 - 5.22<br>2.62 - 2.96 | Baseline<br>Nicotine, 2 mg/kg pre | Baseline<br>Nicotine |
| C57BL (WT) Female<br>n=4 | 12 mo | 22 - 32g<br>24 - 34g | 3.70 - 4.51<br>2.96 - 4.00 | Baseline<br>Nicotine, 2 mg/kg pre | Baseline<br>Nicotine |
| **$\alpha 2$KO Mice** |  |  |  |  |  |
| $\alpha 2$KO Male<br>n=4 | 12-16<br>mo | 22 - 60g<br>18 - 56g | 4.81 - 5.03<br>3.26 - 5.70 | Baseline<br>Nicotine, 2 mg/kg pre | Baseline<br>Nicotine |
| $\alpha 2$KO Female<br>n=4 | 12-16<br>mo | 36 - 42g<br>34 - 38g | 3.70 - 4.40<br>3.66 - 4.66 | Baseline<br>Nicotine, 2 mg/kg pre | Baseline<br>Nicotine |
| **Hypersensitive $\alpha 2$ Mice** |  |  |  |  |  |
| Hypersensitive $\alpha 2$<br>Male n=4 | 12-16<br>mo | 22 - 30g<br>28 - 50g | 4.12 - 4.81<br>4.12 - 4.81 | Baseline<br>Nicotine, 2 mg/kg pre | Baseline<br>Nicotine |
| Hypersensitive $\alpha 2$<br>Female n=4 | 12-16<br>mo | 28 - 50g<br>22 - 28g | 3.92 - 4.51<br>3.66 - 4.22 | Baseline<br>Nicotine, 2 mg/kg pre | Baseline<br>Nicotine |

### 2.2 PET/CT Imaging

Mice were fasted in the scanner room, in a dark quiet place, for 18-24 hours prior to experiments. [^18^F]FDG in sterile saline was purchased commercially from PETNET. On the day of the scan, all mice were placed in an enclosure maintained at 74-78°F. [^18^F]FDG was injected intraperitoneally with 3 to 5 MBq (Table-1) in 0.01 to 0.1 mL 0.9% sterile saline, while awake without isoflurane anesthesia. Subjects were then returned to their cages and allowed to uptake the tracer in an awake, unrestrained state for 40 minutes. The subjects were then anesthetized with isoflurane, 3% induction, maintained at 2.5% and placed inside Siemens Inveon PET/CT scanner. The PET and CT scanners were used in the “docked mode” thus enabling the acquisition of both the PET and CT scans of the mice in the same position for attenuation correction and co-registration. For all the mice, a CT scan (7 minutes) was first acquired followed by a 30-minute static PET scan. After the scan, the mice were returned to their cages. After an interval of at least a week, a second PET/CT scan on the mice were carried out, which included a co-injection of nicotine (intraperitoneally, 2 mg/kg in 0.2 mL sterile saline) along with [^18^F]FDG.

### 2.3 Image Analysis

The PET data was reconstructed as 128×128×159 matrices with a transaxial pixel of 0.776 mm using a OSEM3D algorithm (2 OSEM iterations, 18 MAP iterations and 1.5 target resolution). PET images were corrected for random coincidences, attenuation and scatter. Calibration was conducted to Bq/cc units using a Germanium-68 phantom (cylinder 6 cm diameter), scanned in the Inveon and reconstructed under the same parameters as the mice. The CT projections were acquired with the detector-source assembly rotating over 360 degrees and 720 rotation steps. A projection bin factor of 2 was used in order to increase the signal to noise ratio in the images. The CT images were reconstructed using cone-beam reconstruction with a Shepp filter with cutoff at Nyquist frequency and binning factor of 2 resulting in an image matrix of 512×512×1008 and a voxel size of 0.052 mm. The CT images were spatially transformed to match the reconstructed PET images.

Brain regions and interscapular BAT (IBAT) were analyzed using Inveon Research Workplace software (IRW, Siemens Medical Solutions, Knoxville, TN) using our previously described procedures [31, 32]. Brain regions such as interpeduncular nucleus (IPN) hippocampus (HIP), thalamus (TH), frontal cortex (FC), cerebellum lobes 8 and 9 (CB 8-9), and rest of the cerebellum (CB) were selected. Some of them including IPN and HIP are known to have α2 nAChR expressions. Activity of [^18^F]FDG was normalized using standard uptake value (SUV= ([^18^F]FDG activity in each volume of interest, VOI in kBq/mL)/ (injected dose (in kBq)/ body weight of each animal in g) methods typically used during the use of [^18^F]FDG [32]. The magnitude of BAT [^18^F]FDG was analyzed using VOIs drawn on PET images for interscapular BAT (IBAT). Similarly to previously described methods [24, 25] the VOIs were delineated visually by auto-contouring the [^18^F]FDG activity that was clearly above normal background activity and delineated visually using the CT image.

### 2.4 Statistical Analysis

The SUV values from the PET images were analyzed in Graphpad Prism 10 and Microsoft Excel 16. Student’s t-test were performed with p values <0.05 indicating statistical significance. Error bars are presented as mean ± standard deviation. [^18^F]FDG uptake in the brain and BAT regions in the 3 groups of mice were compared, males and females separately. In addition, [^18^F]FDG uptake in the brain and BAT regions in the 3 groups of mice were compared before and after nicotine treatment.

## 3. Results

### 3.1 Control Mice

Consistent with our previous findings in control mice, both male and female control mice exhibited good brain uptake under baseline conditions in ambient temperatures (Fig 1A and 1C) [32]. Thalamus (TH), interpeduncular nucleus (IPN), frontal cortex (FC), hippocampus (HP) and cerebellum (CB areas 8-9) exhibited higher levels while cerebellum exhibited lower levels (Fig 1A and 1C). Interscapular BAT (IBAT) showed lower uptake under baseline conditions without nicotine. In the presence of nicotine, major changes occurred in brain and IBAT in both male and female mice. A significant increase in the uptake of [^18^F]FDG was seen in the IBAT along with a decrease in uptake in all regions of the brain.

**FIGURE 1:**
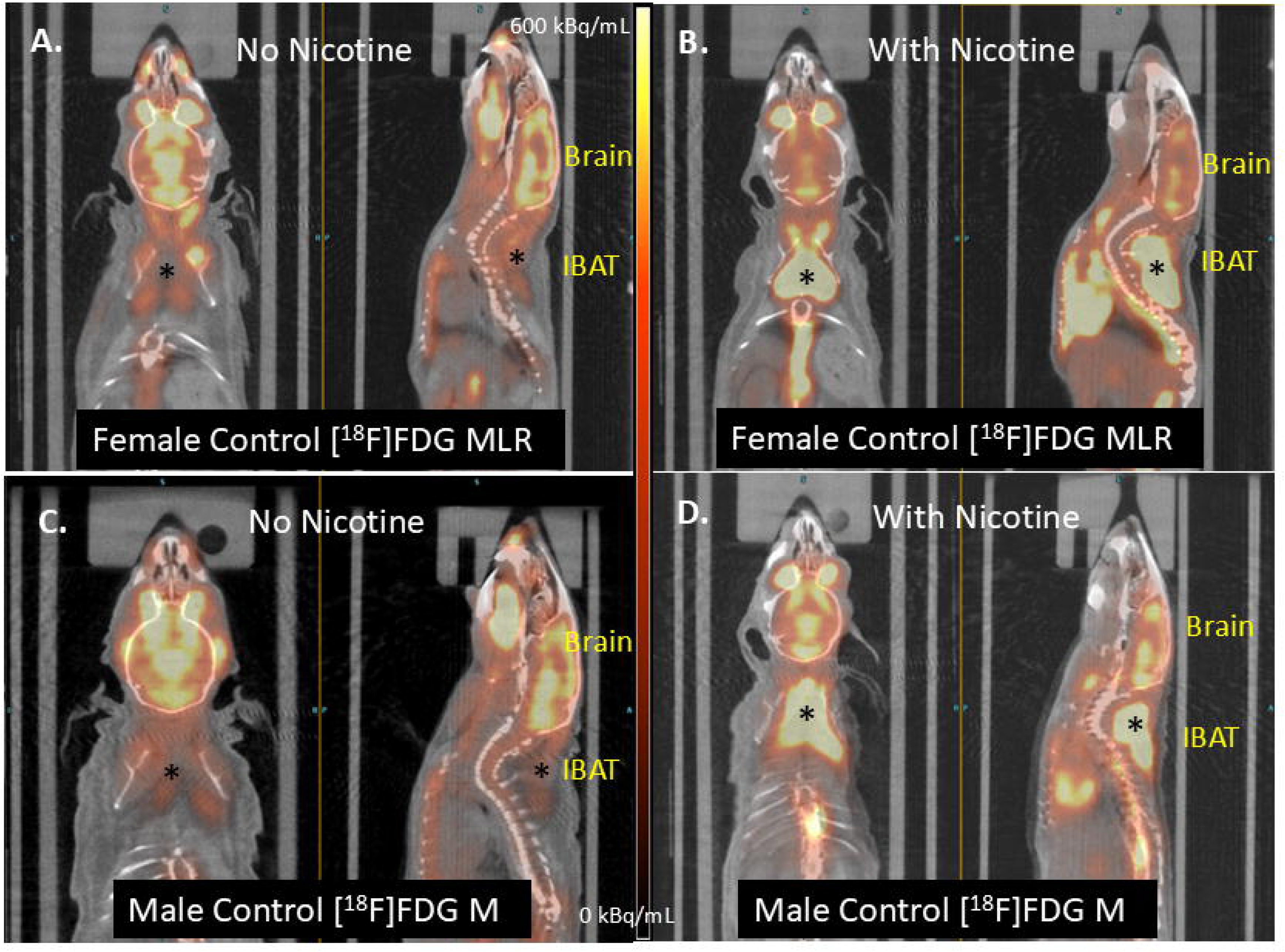
[^18^F]FDG uptake in C57BL/J female mouse (A), same female mouse after pre-injection of nicotine (2 mg/kg) (B), male mouse (C), and same male mouse after pre-injection of nicotine (2 mg/kg) (D). Interscapular brown adipose tissue (IBAT) is marked by an asterisk (*). Nicotine-induced significant activation of IBAT while reducing brain uptake (B,D).

Brain [^18^F]FDG brain uptake ranging approximately from SUV of 3.5 to 5.5 (Fig 2A). Thalamus exhibited higher levels while CB exhibited lower levels (Fig 2A). Female mice exhibited significantly lower average brain uptake compared to male controls by approximately 20% (average females SUV=3.89; average males SUV=4.68; Figure 2B). Nicotine reduced [^18^F]FDG brain uptake in both male and female mice in all the brain regions with SUV levels dropping below 4. Both the nAChR-rich regions (TH) and nAChR-poor regions (CB) were similarly affected (Fig 2A). This global brain reduction of glucose metabolism caused by nicotine in both the male and female mice was highly significant (Fig 2B) Nicotine reduced the [^18^F]FDG SUV average values from 4.68 to 3.38 (28% decrease) in the male mice, while in the female mice it was reduced from 3.89 to 2.53 (35% decrease). The effect of the same dose of nicotine in male and female mice appeared to indicate a greater nicotine-induced [^18^F]FDG reduction in the female mice (Fig 2B).

**FIGURE 2:**
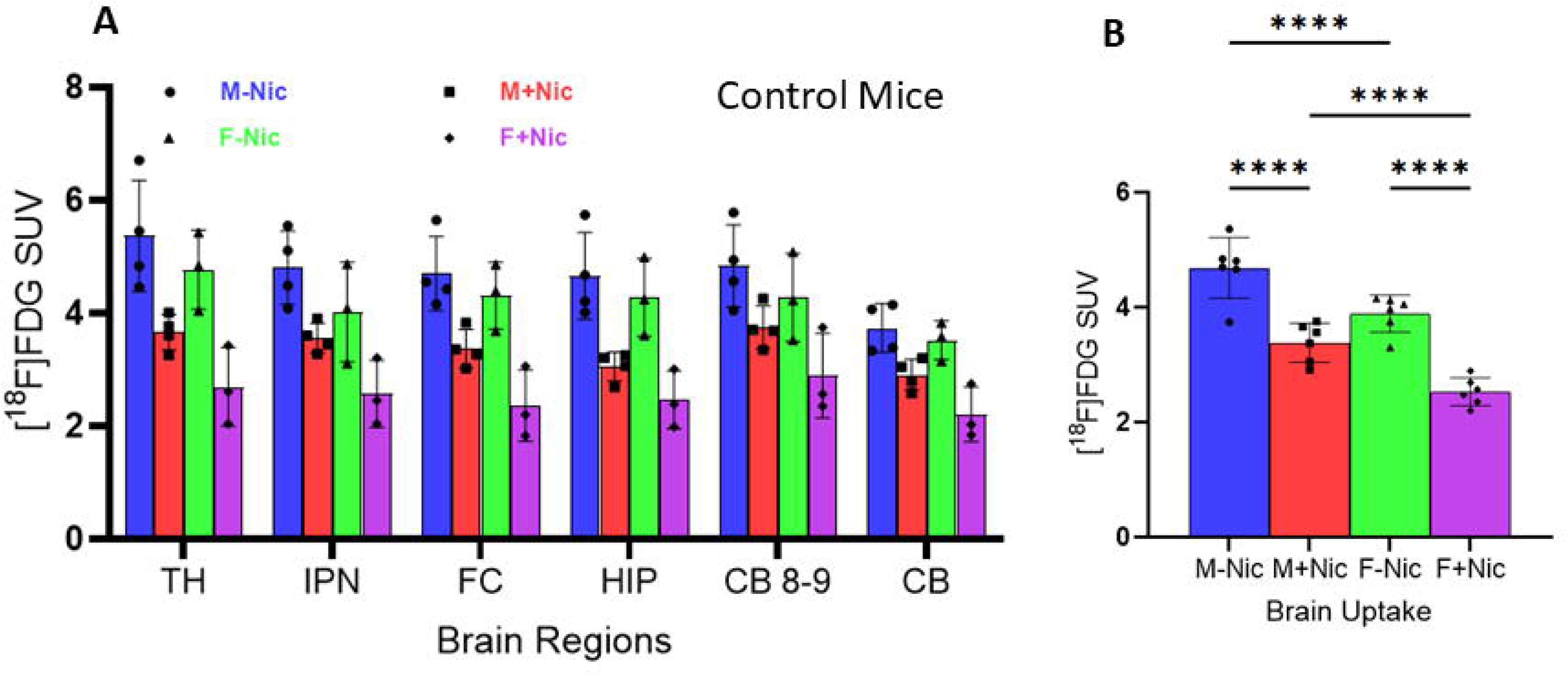
(A). Regional brain distribution of [^18^F]FDG SUV (standard uptake value) in control C57BL/J male and female mice before nicotine (M-Nic and F-Nic) and after nicotine (2 mg/kg, ip; M+Nic and F+Nic) administration (TH: thalamus; IPN: interpeduncular nucleus; FC: frontal cortex; HIP: hippocampus; CB 8-9: cerebellum regions 8 and 9; CB: rest of the cerebellum). (B). Whole brain [^18^F]FDG SUV average of C57BL/J male and female mice showing significant difference (**** p < 0.0001) across sex and nicotine pretreatment. Nicotine pretreatment led to lower [^18^F]FDG SUV in both males and females. Females showed lower [^18^F]FDG SUV compared to males under both conditions.

In the same groups of mice, [^18^F]FDG uptake in IBAT and white adipose tissue (WAT) in the interscapular region were analyzed (Fig 1). Under baseline conditions (no nicotine) male mice exhibited lower IBAT uptake (SUV= 1.74) compared to the brain uptake. However, female mice exhibited significantly higher IBAT uptake (IBAT SUV=2.88) compared to the male mice. As expected, WAT had lower levels of [^18^F]FDG uptake in both males and female mice. Contrary to nicotine treatment reducing [^18^F]FDG uptake in the brain, the uptake of [^18^F]FDG in the IBAT was dramatically increased in the IBAT of both male and female mice. Nicotine increased the IBAT [^18^F]FDG SUV values from 1.74 to 3.97 (a 228% increase) in the male mice, while in the female mice it increased from 2.88 to 6.10 (a 212% increase) (Fig 3). Although the effect of nicotine in male and female mice IBAT appeared similar, the resulting nicotine-induced [^18^F]FDG increase in the female mice IBAT was high (Fig 3). Female mice exhibited significantly higher IBAT uptake compared to nicotine treated male mice by approximately 154% (females SUV=6.10; males SUV=3.97; Figure-3). Thus, the sex effects on [^18^F]FDG uptake are inversed in the IBAT compared to the brain of these control mice. In the case of WAT, the effect of nicotine was statistically insignificant (Fig 3).

**FIGURE 3:**
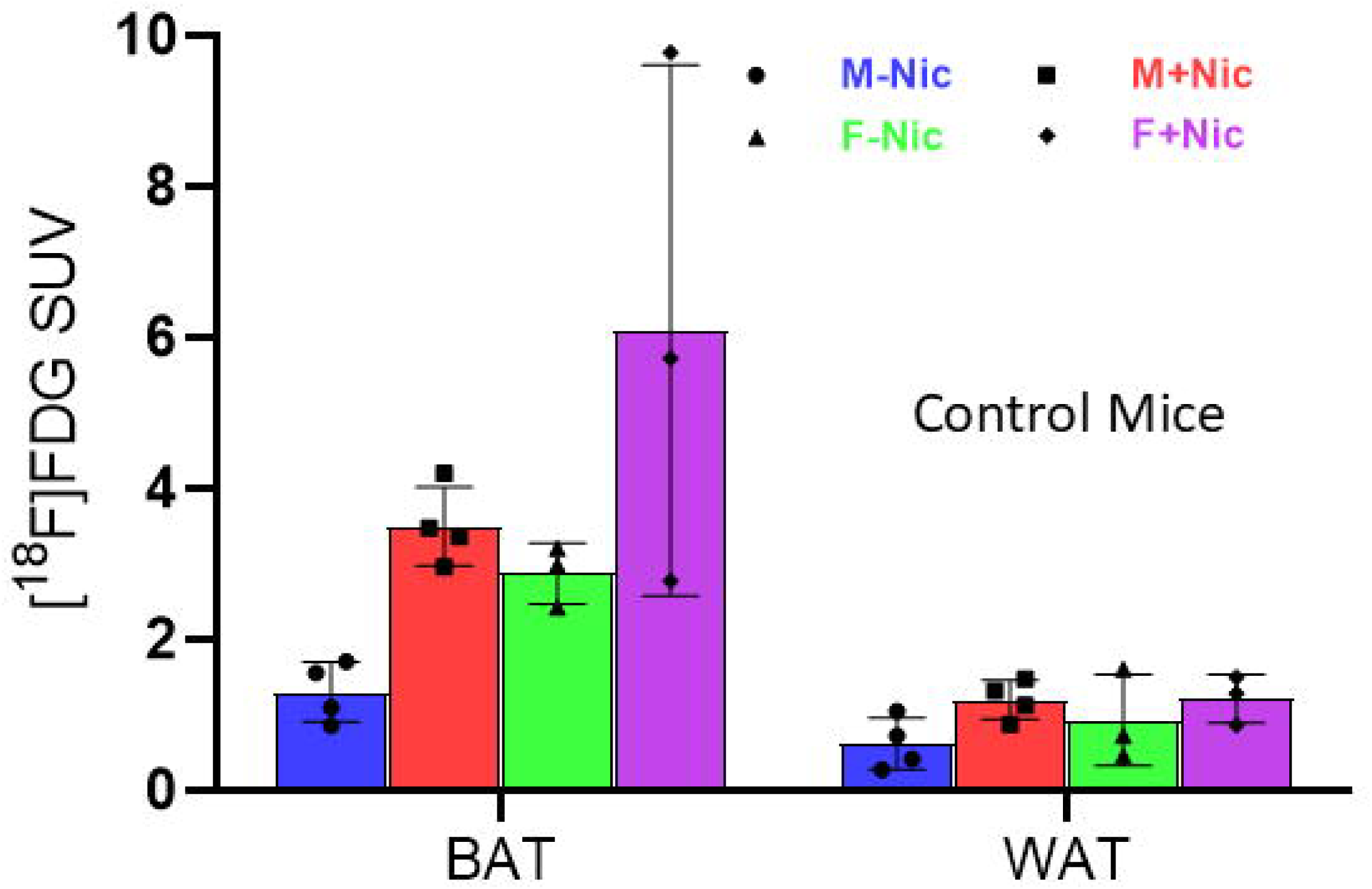
Brown adipose tissue (BAT) and white adipose tissue (WAT) [^18^F]FDG SUV (standard uptake value) in control C57BL/J male and female mice before nicotine (M-Nic and F-Nic) and after nicotine (2 mg/kg, ip; M+Nic and F+Nic). BAT uptake was higher than WAT under all conditions and nicotine pretreatment increased [^18^F]FDG SUV. Females showed higher BAT activity compared to males.

### 3.2 α2 Knock-out Mice

In comparison to the CN mice, major differences of [^18^F]FDG uptake in the male and female α2KO mice were observed (Fig 4). Both male and female α2KO mice exhibited good brain uptake under baseline conditions in ambient temperatures (Fig 4A and 4C). Uptake of [^18^F]FDG was seen in the TH, IPN, FC, HP and CB areas 8-9 similar to the CN mice (Fig 4A and 4C). Interscapular BAT (IBAT) showed very high uptake under baseline conditions without nicotine in both male and female mice (Fig 4A and 4C). In the presence of nicotine, major changes occurred in brain and IBAT in both male and female mice. A significant decrease in the uptake of [^18^F]FDG was seen in the IBAT along with a decrease in uptake in all regions of the brain.

**FIGURE 4:**
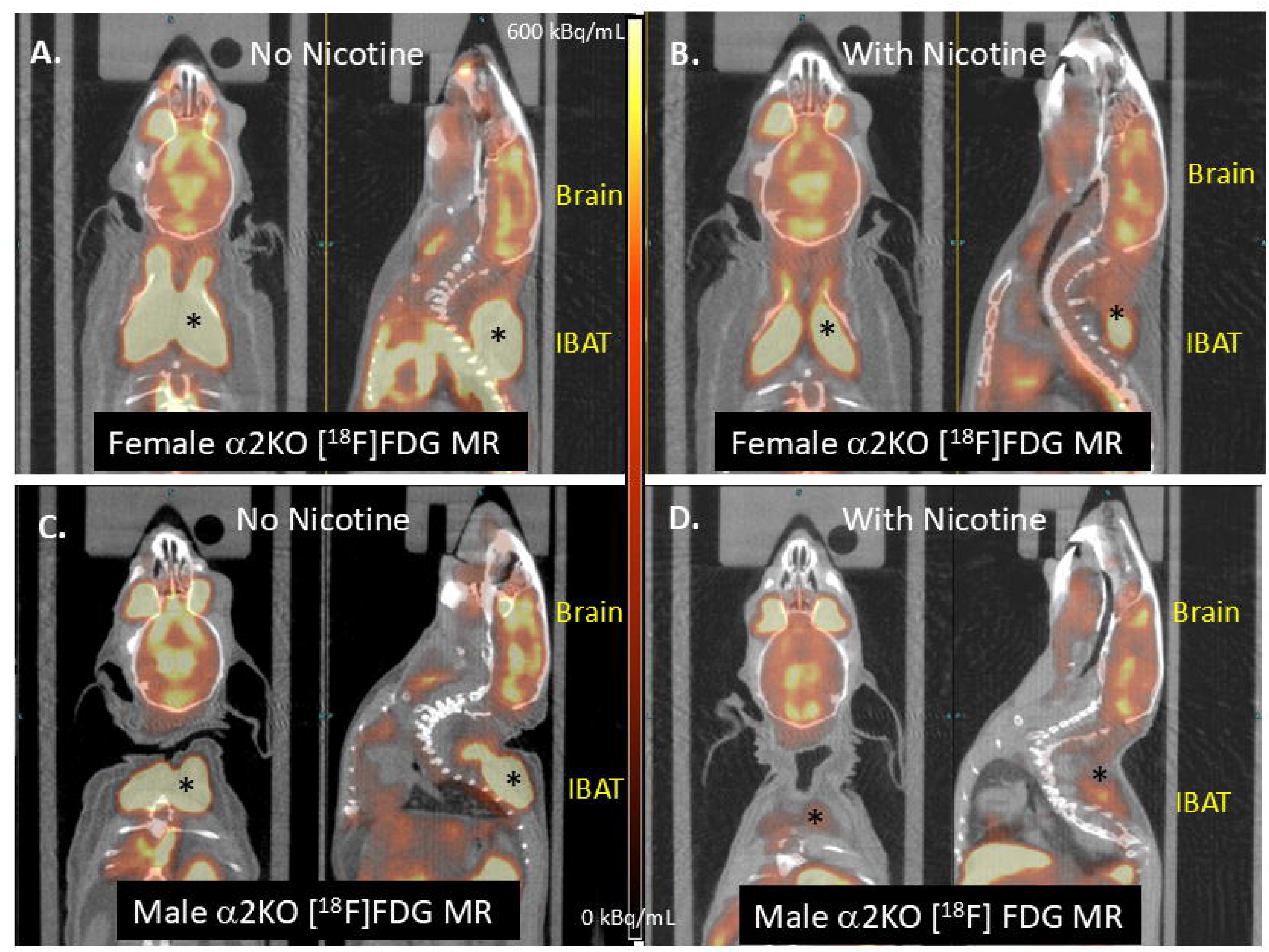
[^18^F]FDG uptake in α2KO female mouse (A), same α2KO female mouse after pre-injection of nicotine (2 mg/kg) (B), male α2KO mouse (C), and same male α2KO mouse after pre-injection of nicotine (2 mg/kg) (D). High [^18^F]FDG uptake in the IBAT (asterisk *) is seen in baseline (A, C). Nicotine-induced significant decrease of IBAT was observed (B,D).

Brain [^18^F]FDG uptake in male and female α2KO mice ranged approximately from SUV of 3.5 to 4.5 (Fig 5A). The uptake was lower compared to the control mice. Female and male mice were similar, with lower discrimination across brain regions compared to the control mice. Female α2KO mice exhibited marginally lower average brain uptake compared to male α2KO mice by approximately 11%, but was significant (females SUV=3.92; males SUV=4.37; Figure 5B). Nicotine reduced [^18^F]FDG brain uptake in both male and female α2KO mice in all the brain regions with SUV levels dropping below 3.5. In female α2KO mice, the reduction in [^18^F]FDG brain uptake was less than that found in the male α2KO mice. This sex effect was seen in both the nAChR-rich regions TH and nAChR-poor regions CB (Fig 5A). The global brain reduction of glucose metabolism caused by nicotine in both the male and female mice was highly significant (Figure 5B) Nicotine reduced the [^18^F]FDG SUV values from 4.37 to 2.98 (a 32% decrease) in the male mice, while in the female mice it was reduced from 3.92 to 3.38 (a 14% decrease). The effect of nicotine in male and female mice appeared to indicate a greater nicotine-induced [^18^F]FDG reduction in the male mice (Figure 5B). This nicotine effect on α2KO mice was the inverse of the findings in the control mice, where the female mice exhibited a greater effect of nicotine. Thus, it appears that the α2 nAChR mediated nicotine effects may be more prominent in female mice compared to male mice.

**FIGURE 5:**
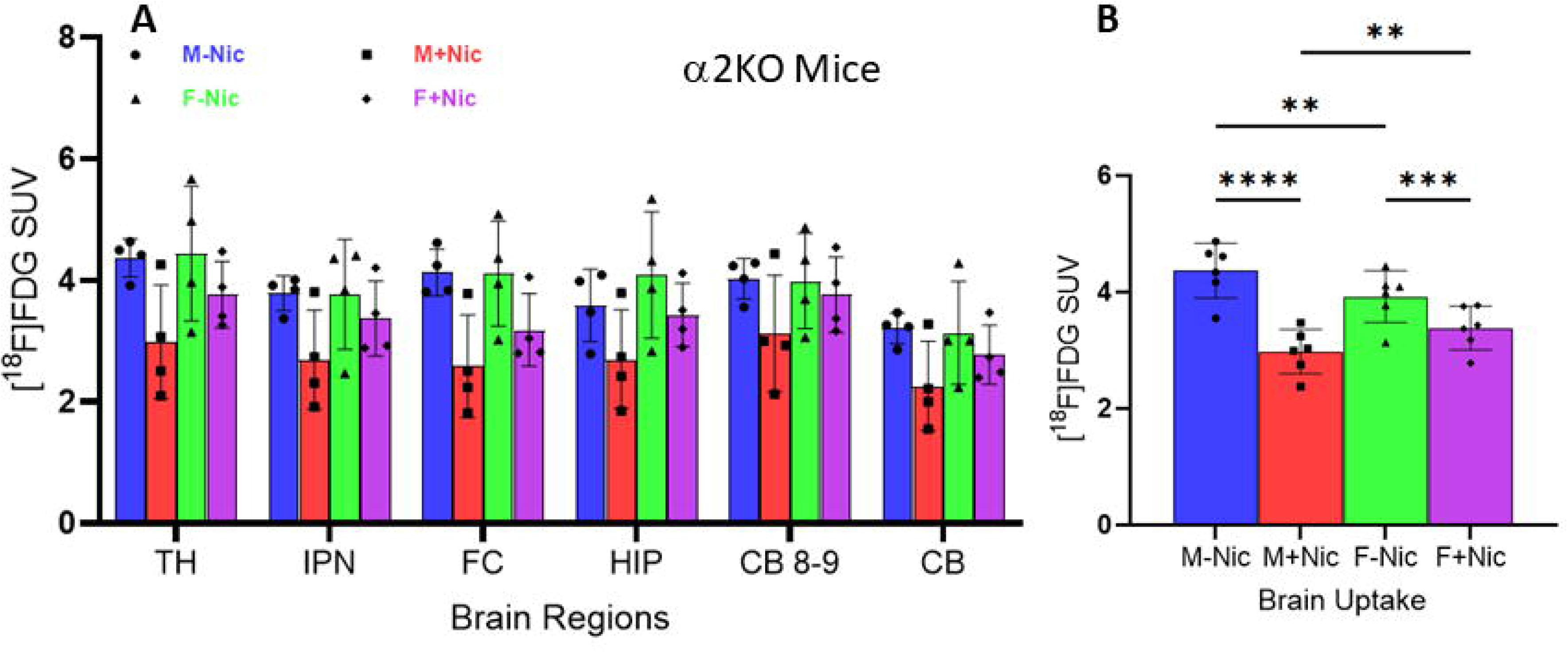
(A). Regional brain distribution of [^18^F]FDG SUV (standard uptake value) in α2KO male and female mice before nicotine (M-Nic and F-Nic) and after nicotine (2 mg/kg, ip; M+Nic and F+Nic) administration (TH: thalamus; IPN: interpeduncular nucleus; FC: frontal cortex; HIP: hippocampus; CB 8-9: cerebellum regions 8 and 9; CB: rest of the cerebellum). (B). Whole brain [^18^F]FDG SUV average of α2KO male and female mice showing significant difference (** p < 0.01, *** p < 0.001, **** p < 0.0001, ns = not significant) across sex and nicotine pretreatment. Nicotine pretreatment led to lower [^18^F]FDG SUV in both males and females. Males showed lower [^18^F]FDG SUV compared to females with nicotine pretreatment.

Analysis of [^18^F]FDG uptake in the IBAT and WAT in the α2KO mice was carried out similar to the control mice (Fig 6). Under baseline conditions (no nicotine), both male and female α2KO mice exhibited unusually high IBAT uptake (SUV= 9.05 for males; SUV= 7.86 for females) compared to the brain uptake. There was little sex effect in both the brain and IBAT in the α2KO mice compared to the control mice. Like the control mice, WAT had lower levels of [^18^F]FDG uptake in the α2KO mice. Similar to nicotine treatment reducing [^18^F]FDG uptake in the brain, the uptake of [^18^F]FDG in the IBAT was dramatically decreased in the IBAT of both male and female α2KO mice (Fig 6). Nicotine decreased the [^18^F]FDG SUV values from 9.05 to 4.37 (a 52% decrease) in the male mice, while in the female mice it decreased from 7.86 to 4.52 (a 42% decrease). This nicotine effect in the α2KO mice was the opposite of the nicotine effect observed in the control mice. Thus, [^18^F]FDG uptake in the IBAT in the α2KO mice compared to the control mice was different in three respects: (1). A very high IBAT [^18^F]FDG uptake was observed in the baseline studies in both male and female mice; (2). Nicotine decreased IBAT [^18^F]FDG uptake in both males and females; (3). There was no significant sex effect between males and females either in the baseline or the nicotine treatment study.

**FIGURE 6:**
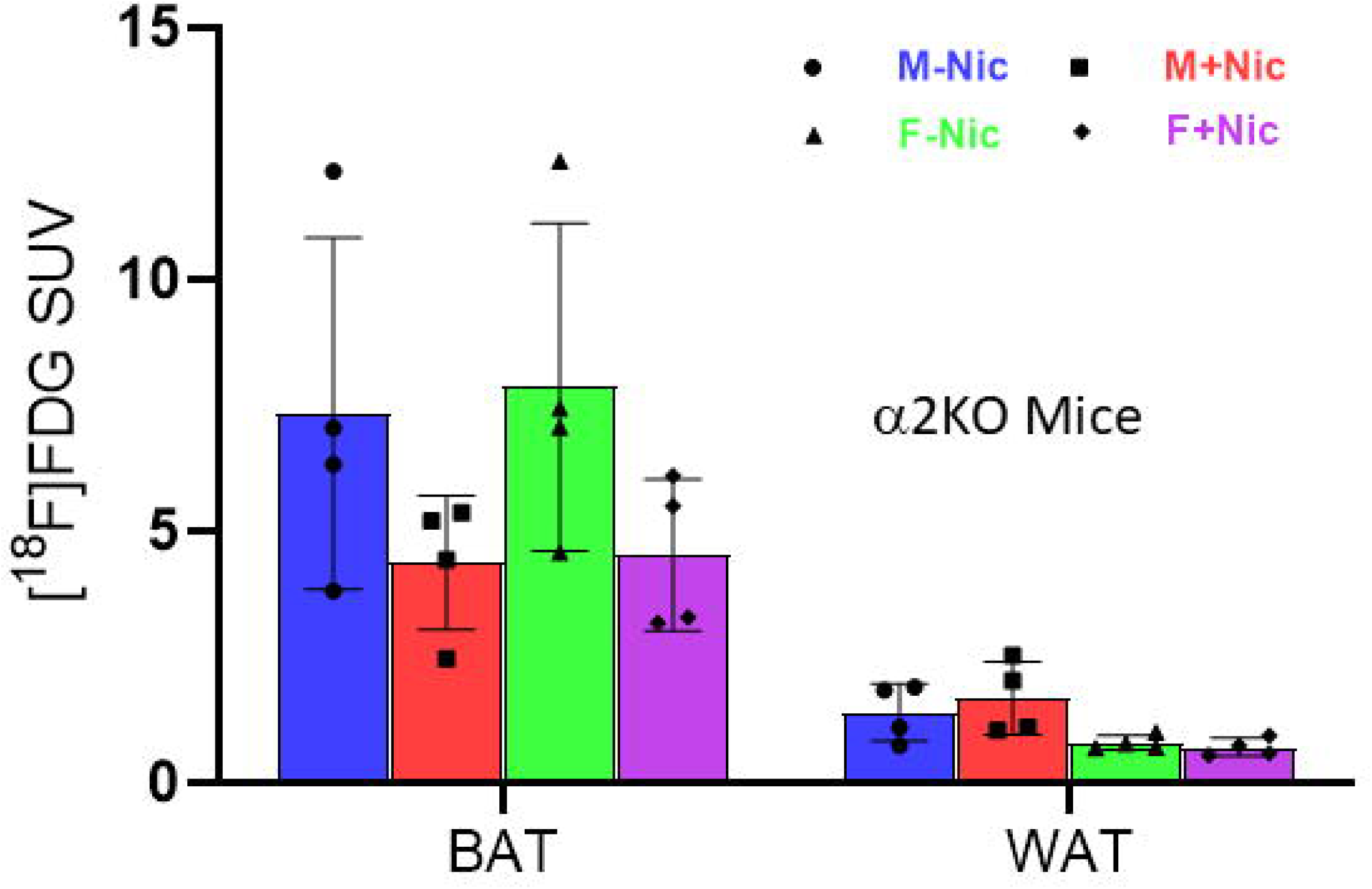
Brown adipose tissue (BAT) and white adipose tissue (WAT) [^18^F]FDG SUV (standard uptake value) in α2KO male and female mice before nicotine (M-Nic and F-Nic) and after nicotine (2 mg/kg, ip; M+Nic and F+Nic). BAT uptake was higher than WAT under all conditions. Nicotine pretreatment decreased [^18^F]FDG SUV in both males and females, with no significant difference between the sexes.

### 3.3 α2 Hypersensitive Mice

The α2HS mice exhibited [^18^F]FDG uptake more similar to the control mice (Fig 7)., Both male and female control mice exhibited good brain uptake under baseline conditions in the various brain regions. However, noticeably, male α2HS mice had significantly higher [^18^F]FDG uptake compared to the female α2HS mice (Fig 7A and 7C). Interscapular BAT (IBAT) showed low uptake under baseline conditions without nicotine similar to the control mice. In the presence of nicotine, a significant increase in the uptake of [^18^F]FDG was seen in the IBAT, while a decrease in uptake in all brain regions was evident.

**FIGURE 7:**
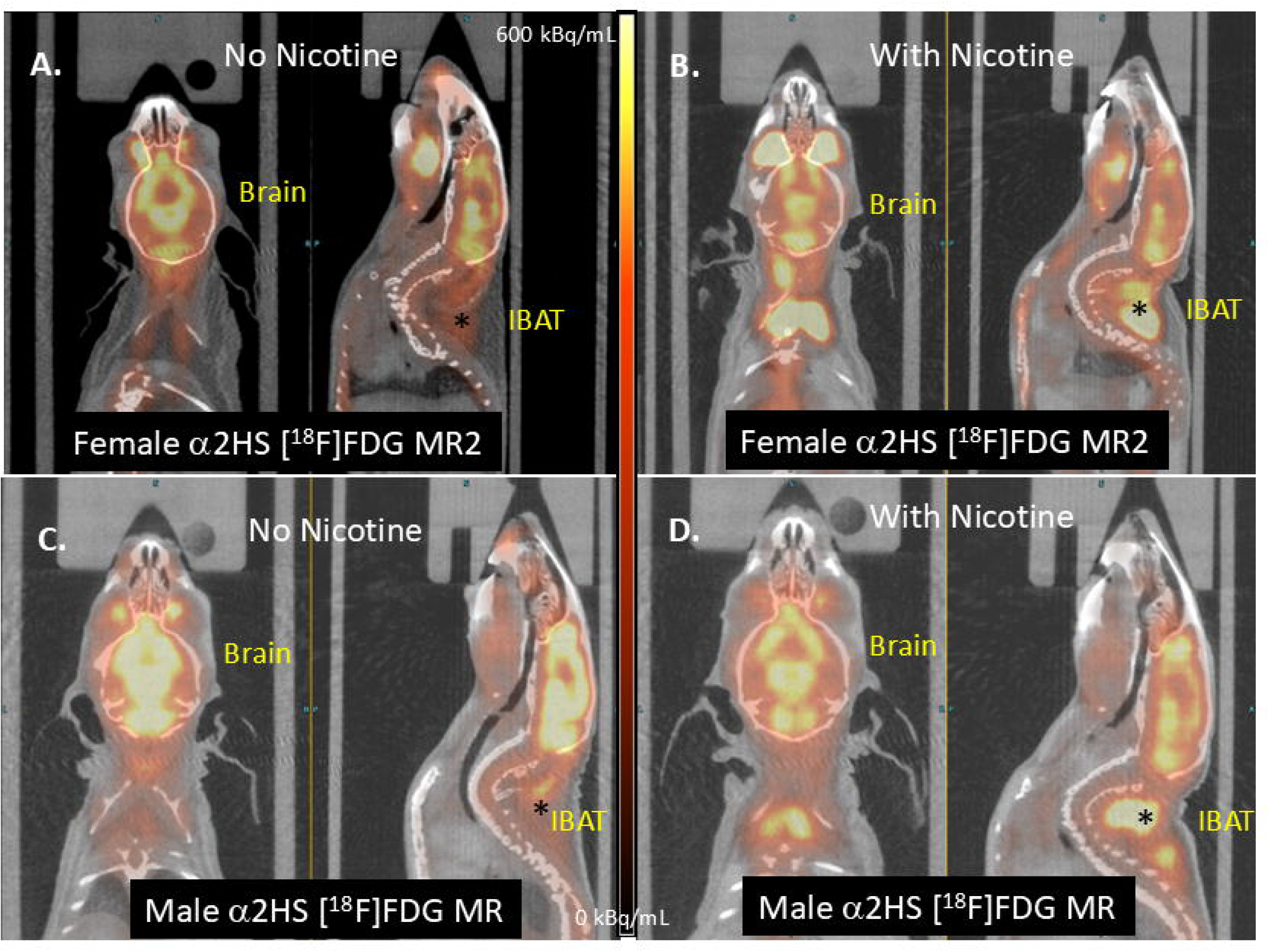
[^18^F]FDG uptake in α2HS female mouse (A), same female α2HS mouse after pre-injection of nicotine (2 mg/kg) (B), male α2HS mouse (C), and same male α2HS mouse after pre-injection of nicotine (2 mg/kg) (D). Interscapular brown adipose tissue (IBAT) is marked by an asterisk (*). Nicotine-induced significant activation of IBAT while reducing brain uptake (B,D).

Although both male and female α2HS mice exhibited good [^18^F]FDG brain uptake, the male α2HS mice exhibited much higher uptake (SUV=>6) versus female α2HS mice (SUV=3.5 to 4). As in the control mice, thalamus exhibited higher levels while cerebellum exhibited lower levels (Fig 8A). Thus, of all the mice, the male α2HS mice exhibited the highest [^18^F]FDG uptake in the brain. Male-female difference in the brain was highly significant. Nicotine treatment caused a significant decrease in [^18^F]FDG uptake in both male and female α2HS mice. The decrease in the male α2HS mice from baseline to nicotine treatment was much larger, approximately 44% (average SUV=5.88 to average SUV=3.30), whereas female mice change was smaller (32%; average SUV=3.63 to average SUV=2.47), but significant (Fig 8B). Thus, it appears that the α2HS nAChR and nicotine may have different mechanisms/pathways that affect glucose metabolism in males and females, with females being less responsive to the increased sensitivity of the α2HS nAChR.

**FIGURE 8:**
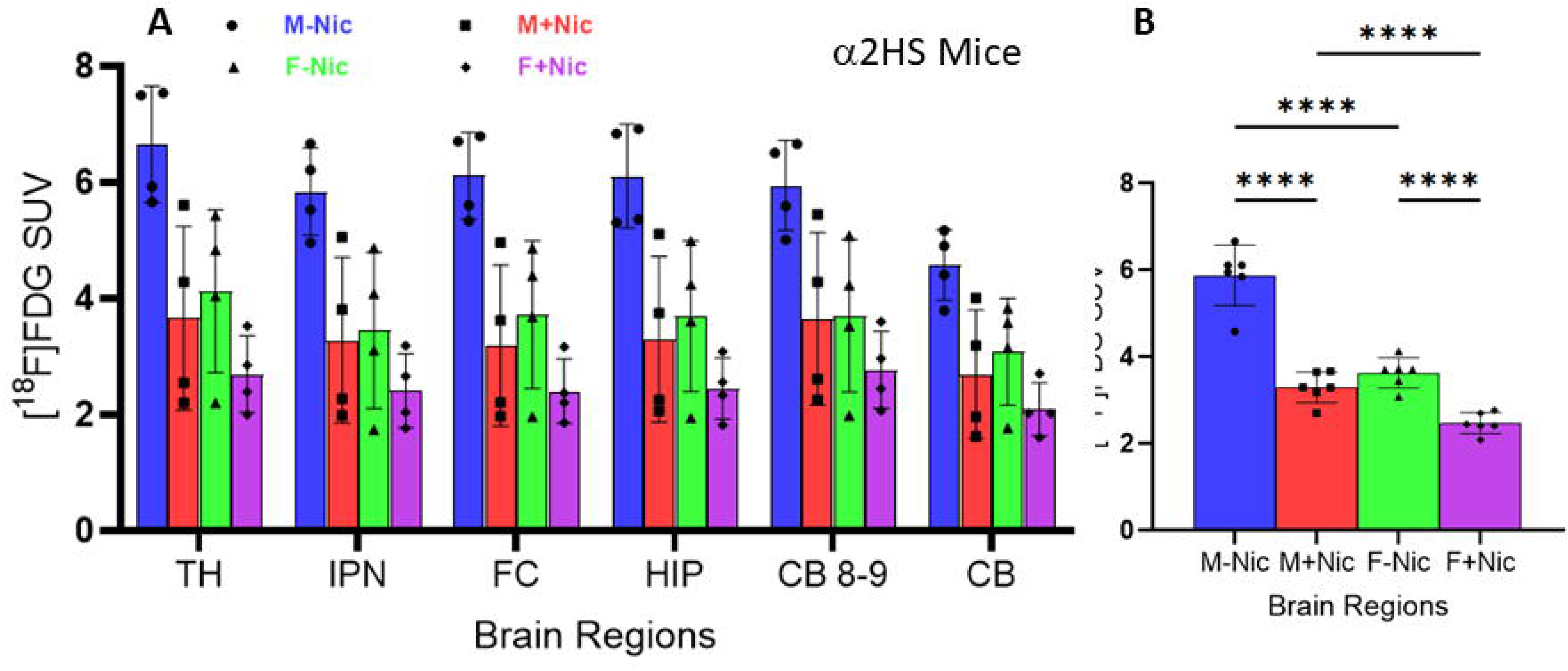
(A). Regional brain distribution of [^18^F]FDG SUV (standard uptake value) in α2HS male and female mice before nicotine (M-Nic and F-Nic) and after nicotine (2 mg/kg, ip; M+Nic and F+Nic) administration (TH: thalamus; IPN: interpeduncular nucleus; FC: frontal cortex; HIP: hippocampus; CB 8-9: cerebellum regions 8 and 9; CB: rest of the cerebellum). (B). Whole brain [^18^F]FDG SUV average of α2HS male and female mice showing significant difference (**** p < 0.0001) across sex and nicotine pretreatment. Nicotine pretreatment led to lower [^18^F]FDG SUV in both males and females. Females showed lower [^18^F]FDG SUV compared to males under both conditions. Males showed a greater decrease with nicotine pretreatment.

[^18^F]FDG uptake in α2HS IBAT and WAT in the interscapular region were similarly analyzed (Fig 9). Although the uptake profile in the two groups of mice under the two conditions were somewhat similar to the control group, there were observable differences. Under baseline conditions (no nicotine) male mice exhibited lower IBAT uptake (SUV= 3.35) compared to the brain uptake. However, female mice exhibited higher IBAT uptake (IBAT SUV=4.04 vs brain SUV=3.63) compared to the male mice. As expected, WAT had lower levels of [^18^F]FDG uptake. The uptake of [^18^F]FDG in the IBAT was dramatically increased in the presence of nicotine for both male and female α2HS mice. This is similar to the control mice but opposite of the α2KO group of mice. Nicotine increased the [^18^F]FDG SUV values from 3.35 to 4.79 (a 143% increase) in the male mice, while in the female mice it increased from 4.04 to 6.02 (a 149% increase). There was no decipherable effect of nicotine on WAT (Fig 9). Nicotine-induced effects appeared to be greater in the female mice (Fig 9). Thus, the sex effects on [^18^F]FDG uptake in the α2HS mice appeared to be similar to the control mice.

**FIGURE 9:**
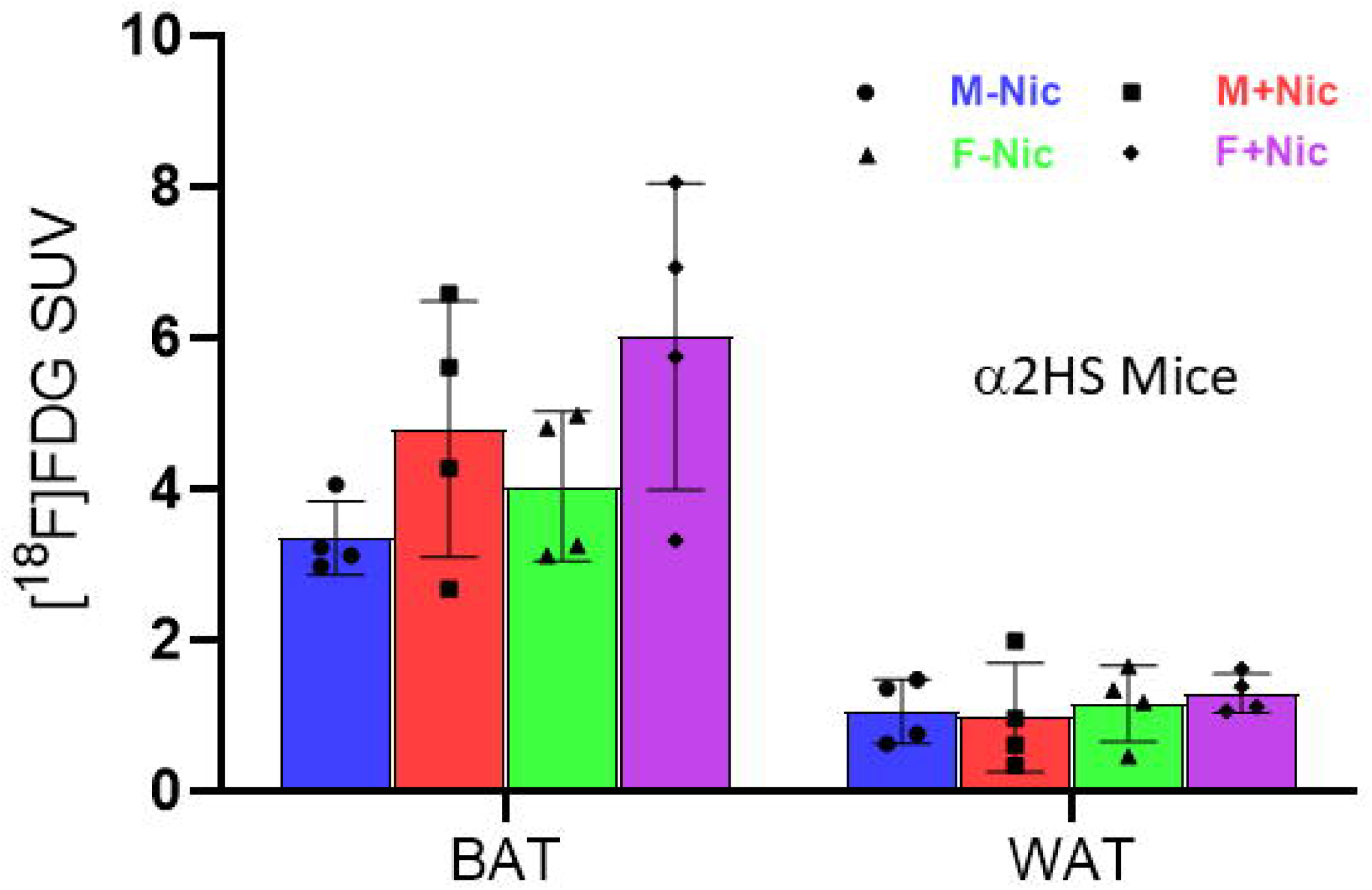
Brown adipose tissue (BAT) and white adipose tissue (WAT) [^18^F]FDG SUV (standard uptake value) in α2HS male and female mice before nicotine (M-Nic and F-Nic) and after nicotine (2 mg/kg, ip; M+Nic and F+Nic). BAT uptake was higher than WAT under all conditions. Nicotine pretreatment increased [^18^F]FDG SUV in both males and females, with females showing higher BAT activity.

### 3.4 Comparing Brain [^18^F]FDG in control, α2KO and α2HS Mice

Comparison of the whole brain [^18^F]FDG uptake in the 3 groups of mice is shown in Fig 10. Male α2HS mice exhibited the significantly higher uptake of [^18^F]FDG compared to control male mice while female α2HS mice did not exhibit a higher uptake compared to control female mice (Figure 10A). Nicotine decreased [^18^F]FDG uptake in all the mice and there was no difference between the α2HS and control male and female mice. Uptake of [^18^F]FDG in the α2KO mice, males and females, were lower to controls without nicotine and were statistically significant. In the presence of nicotine, there was a reduction of [^18^F]FDG. However, female α2KO mice exhibited a lower reduction compared to the male α2KO mice and the control female mice (Fig 10B).

**FIGURE 10:**
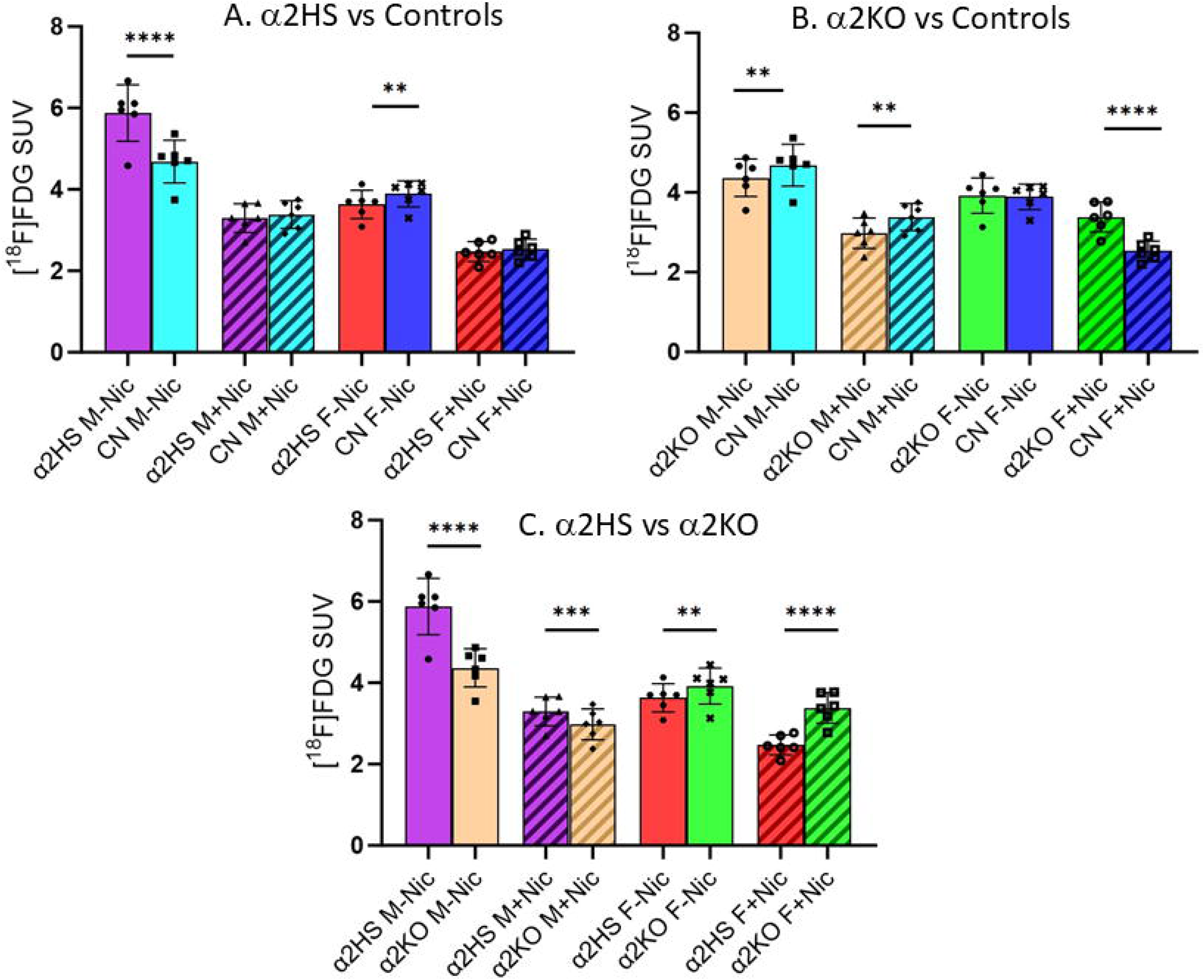
Brain comparison of brain [^18^F]FDG SUV in the three groups of mice. Comparison of brown adipose tissue (BAT) [^18^F]FDG SUV (standard uptake value) in α2HS, control C57BL/J and α2KO male and female mice before nicotine and after nicotine (-NIC and +NIC; 2 mg/kg, ip). Without nicotine, α2KO male and female mice showed higher BAT activity compared to the α2HS and control C57BL/J mice. Nicotine pretreatment increased [^18^F]FDG SUV in both males and females of α2HS and control C57BL/J, but decreased BAT activity of α2KO male and female mice. (** p < 0.01, *** p < 0.001, **** p < 0.0001).

The α2HS male mice exhibited the largest difference in [^18^F]FDG compared to α2KO male mice in the absence of nicotine. The female α2HS mice did not exhibit this large a difference compared to α2KO female mice, although both males and females were statistically significant. In the presence of nicotine, there was a similar reduction of [^18^F]FDG in the male α2HS and α2KO mice. Female α2HS mice exhibited a significantly larger reduction compared to the female α2KO mice (Fig-10C). Thus, the rank order of brain [^18^F]FDG uptake in the 3 groups of mice was: α2HS♂> CN♂> α2KO♂> CN♀= α2KO♀≥ α2HS♀. In the presence of nicotine, the rank order changed to: α2HS♂= CN♂≥ α2KO♂> = α2KO♀> α2HS♀= CN♀. Females had lower [^18^F]FDG uptake compared to males, with the only exception of α2KO females had higher uptake compared to α2KO males in the presence of nicotine.

### 3.5 Comparing IBAT [^18^F]FDG in control, α2KO and α2HS Mice

Figure 11 shows comparison of [^18^F]FDG uptake in IBAT in the 3 groups of mice in the absence and presence of nicotine. The major difference was the IBAT of α2KO male and female mice compared to CN and α2HS. The α2KO mice had significantly higher baseline [^18^F]FDG uptake compared to the other two groups. More importantly, nicotine decreased IBAT [^18^F]FDG in the α2KO mice rather than the expected increases seen in the CN and α2HS mice. However, the levels of [^18^F]FDG uptake in the α2KO mice were higher compared to the baseline levels of CN and α2HS mice. Thus, the rank order of IBAT [^18^F]FDG baseline uptake in the 3 groups of mice was: α2KO♂> α2KO♀> α2HS♀> α2HS♂> CN♀> CN♂. Females had higher [^18^F]FDG uptake compared to males, with the only exception of α2KO females had lower uptake compared to α2KO males at baseline. In the presence of nicotine, the rank order changed to: α2HS♀= CN♀ > α2HS♂≥ α2KO♂> = α2KO♀> CN♂.

**FIGURE 11:**
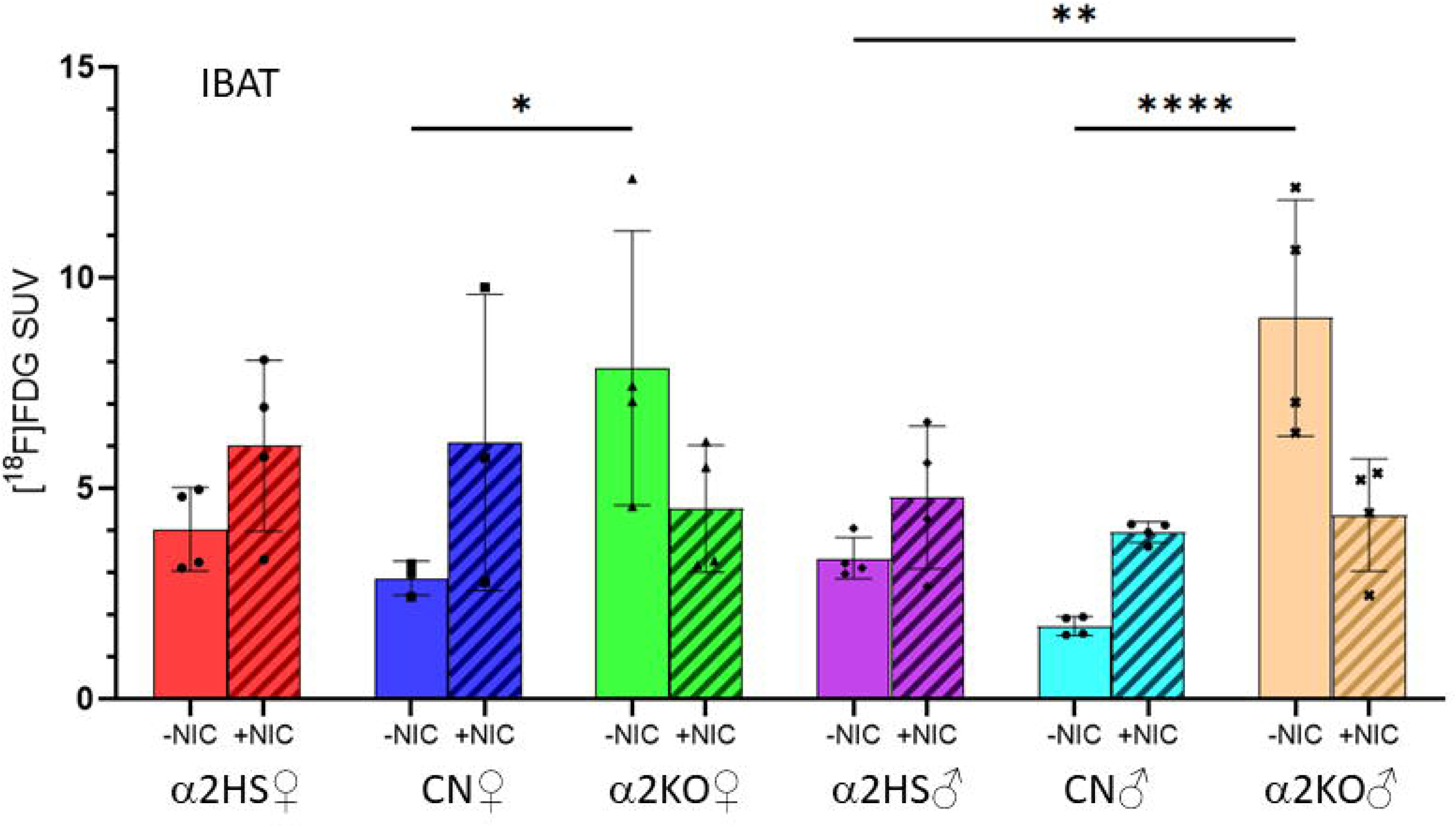
IBAT comparison of [^18^F]FDG SUV (standard uptake value) in α2HS, control C57BL/J and α2KO male and female mice before nicotine and after nicotine (-NIC and +NIC; 2 mg/kg, ip). Without nicotine, α2KO male and female mice showed higher BAT activity compared to the α2HS and control C57BL/J mice. Nicotine pretreatment increased [^18^F]FDG SUV in both males and females of α2HS and control C57BL/J, but decreased BAT activity of α2KO male and female mice. (* *p* < 0.05, ** p < 0.01, **** p < 0.0001).

## 4. Discussion

Although low in abundance, the α2 nAChRs in the human and monkey brain possess a higher affinity for nicotine compared to the other subtypes [<u>6</u>, <u>7</u>). These α2 nAChRs play a critical role in cortical function, mediate response to nicotine, play a role in plasticity and fear learning and memory that is modulated by nicotine [5, 9, 33]. In the sympathetic nervous system which controls adipose activity, BAT and beige-adipocytes have been shown to be abundant in α2 nAChRs [27, 28]. In this study we have used the α2KO and α2HS mice to shed additional light on functional aspects of this receptor subtype.

[^18^F]FDG PET measures glucose metabolic activity [34], and it serves as a valuable non-invasive imaging tool for studies related to synaptic neurotransmission to study brain functional changes requiring glucose [35]. Our goal was to study the potential role of the α2 nAChR mediated neurotransmission on glucose metabolism. Previous reports in rodents [36, 37] and humans have described the male-female differences in [^18^F]FDG brain uptake [38, 39] and cognitive reserve with aging [40]. In the present study, male-female differences in [^18^F]FDG brain uptake were observed. Male mice showed a higher uptake compared to female mice and α2HS mice exhibited the highest brain uptake of [^18^F]FDG. Uptake of [^18^F]FDG in the α2KO mice, males and females, were lower compared to controls and were statistically significant, suggesting a role of α2 nAChR in neurotransmission. This was further supported by the observation that α2HS male mice exhibited the largest increase in [^18^F]FDG compared to α2KO and control male mice. Surprisingly, the female α2HS mice did not exhibit such increased glucose metabolic activity compared to the control and α2KO female mice. Overall, females had lower [^18^F]FDG uptake compared to males and were less sensitive to the presence or absence of α2 nAChR.

A number of human and nonhuman studies have used [^18^F]FDG PET to study effects of nicotine exposure on gene alterations [41] and efforts related to tobacco and smoking use [42, 43]. The action of nicotine on several nAChR subtypes directly, and indirectly on other CNS neurotransmitter-receptor systems contributes to the overall reduction in glucose metabolic activity, suggesting a decrease in synaptic neurotransmission. In this study nicotine had a similar effect of reducing glucose metabolic activity in all the 3 groups. The α2KO mice appeared to be less sensitive to nicotine compared to the control and α2HS mice. This effect was prominent in the α2KO female mice. The effect of nicotine was greatest in the α2HS males because of their higher baseline metabolic activity. Nicotine causes a larger decrease in metabolic (neurotransmitter activity) in males, compared to females and this may cause the sex differences in fear conditioning in female compared to male mice [9]. The enzyme aromatase which is closely linked to estrogen levels is associated with maintaining glucose metabolism and sex hormones play a major role [44, 45]. Nicotine is a known inhibitor of aromatase [46, 47] and was additionally confirmed using PET experiments in female human subjects [48]. The female α2KO mice were more resistant to the nicotine-induced reduction in glucose metabolism compared to the α2HS and control mice. This suggests that the α2 nAChR plays a significant role in nicotine effects in the female mice.

Brown adipose tissue (BAT) is well known as a major glucose disposal tissue [49]. It is known to be a major factor in energy homeostasis as well as being responsible for thermogenesis, and the adipocytes that make up the brown adipose tissue secrete factors that manage insulin, body weight, and inflammation [50]. The α2 subtype is abundant within the brown and beige adipose tissue [27] and has been found to have a role in increased obesity in response to high-fat diet and related metabolic disorders [28]. Nicotine and tobacco can increase thermogenesis in BAT [43, 44]. Nicotine can induce the browning of WAT through the κ opioid receptor [51] and lower WAT weight and significantly increase BAT cell count dependent on nicotine dose [52].

In the absence of regulatory control by the α2 nAChR, the α2KO mice IBAT, both males and females exhibited significantly higher [^18^F]FDG IBAT uptake compared to the controls and α2HS mice. The high level of [^18^F]FDG uptake in the α2KO mice IBAT is unusual compared to the reported several other mice model studies [53]. This suggests that the α2 nAChR has a complex role on IBAT glucose metabolism. Thus, non-α2 nAChR pathways may be involved in IBAT activation of α2KO mice [<u>54</u>, <u>55</u>). Additionally, effects of central nervous system on the sympathetic nervous system of α2KO mice may influence glucose metabolic pathways [56]. At baseline (no nicotine), the hypersensitive, α2HS mice IBAT were lower compared to α2KO mice but still in the range of control mice IBAT.

In the case of nicotine treatment, male and female mice in the 3 groups behaved similarly with increased amount of IBAT activity compared to the brain in all groups. Nicotine induced a reduction in glucose metabolism in the α2KO mice, while in the control and α2HS mice, there was an increase in glucose metabolism which was consistent with the effects of nicotine in control IBAT [21, 22]. The nicotine-induced reduction of IBAT [^18^F]FDG in the α2KO mice was surprising and suggests a complex pathway of nicotine-induced BAT activity. The hypersensitive, α2HS mice IBAT exhibited the greatest increase in IBAT [^18^F]FDG which may be consistent with the higher amount of α2 nAChRs located in beige adipocytes, which are known to take up glucose [28].

Overall, our findings are consistent with these previous studies, showing reduced brain activity across all three groups (with a lesser effect on the α2KO mice). The α2KO mice also showed a reduction in activity in the IBAT after the addition of nicotine, suggesting that without these receptors, nicotine has the opposite relationship with these tissues. Chronic tobacco use was also significantly associated with decreased hepatic metabolic activity as measured on [^18^F]FDG-PET/CT, which suggests that smoking induces an adverse hepatotoxic effect. Nicotine abuse was also associated with decreased hepatic metabolic activity, suggesting lowered [^18^F]FDG (glucose) metabolism [42]. Figure 12 shows a summary of our findings and the general trend of [^18^F]FDG distribution in the brain and IBAT across our three groups and with the addition of nicotine. α2 nAChRs have a significant effect on [^18^F]FDG uptake in the brain in male hypersensitive mice. However, for all other groups, the effect in the brain is minimal. The addition of nicotine led to a decrease in brain uptake across all three groups regardless of sex. When it comes to IBAT [^18^F]FDG uptake, there is a significant effect of nicotine on all three groups and both males and females, but findings of α2KO mice need further studies.

**FIGURE 12:**
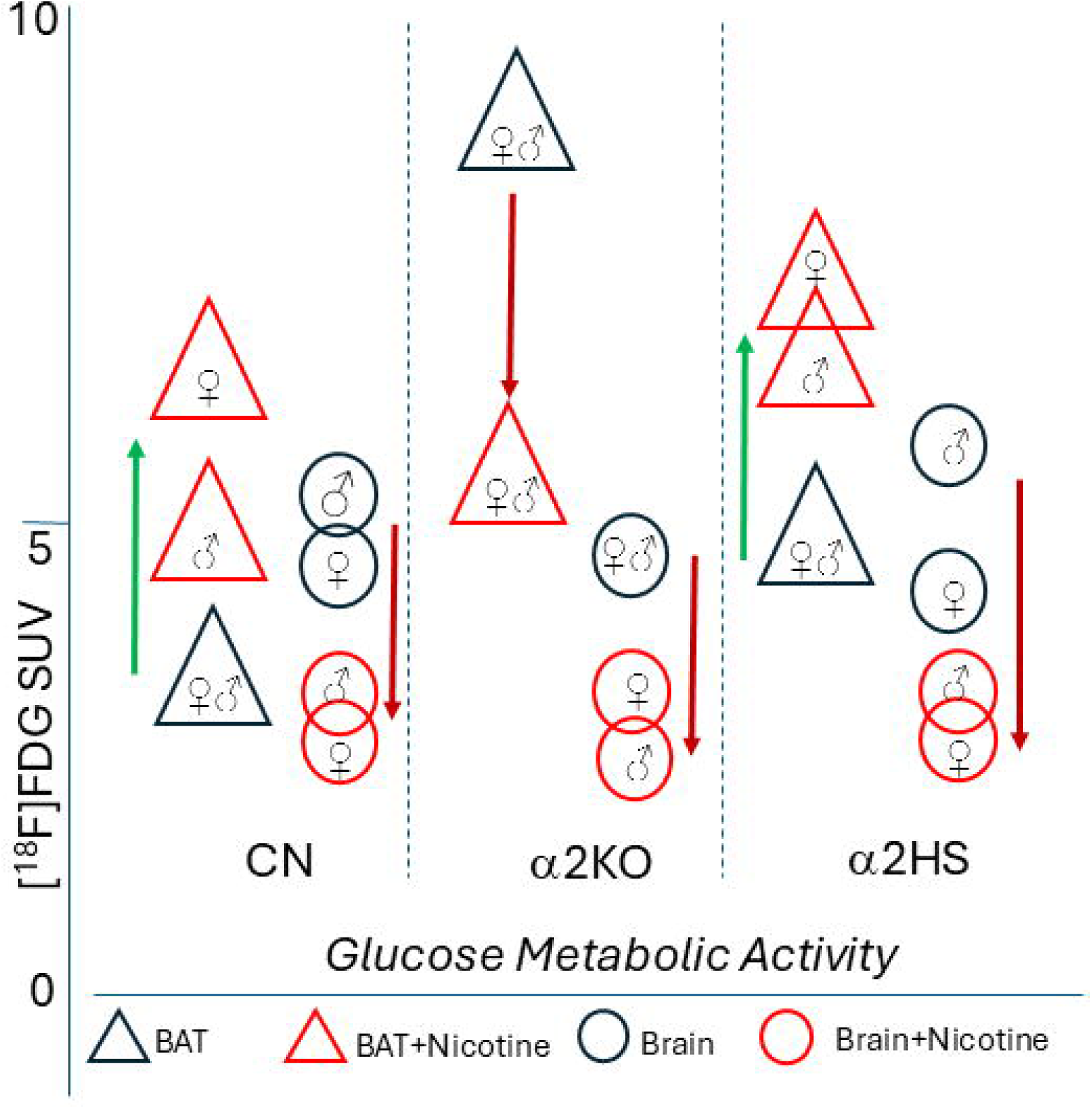
Summary of brain and BAT [^18^F]FDG SUV in the three groups of mice.

In our assessment, following are some of the limitations of this study: 1. All [^18^F]FDG imaging studies were done using the standard protocol of fasting the mice to maximize brain uptake. Future studies may consider non-fasted mice to compare brain and IBAT uptake; 2. Use of lower doses of nicotine may be useful to compare the 3 groups and provide further information on the α2HS mice; (3). Although significant differences amongst the 3 groups of animals were deciphered, Increasing the number of animals would further strengthen the data.

## 5. Conclusions

Brain uptake of [^18^F]FDG in the α2KO mice, males and females, were lower compared to controls and higher in α2HS mice suggesting a role of α2 nAChR in neurotransmission. Male-female differences were observed, with females having lower brain [^18^F]FDG uptake compared to males. Females were less sensitive to the presence or absence of α2 nAChR. Brain uptake in female mice was also less responsive to nicotine compared to the male mice.

[^18^F]FDG uptake in the IBAT in α2KO mice compared to the control mice was different in three respects: (1). A very high IBAT [^18^F]FDG uptake was observed in the baseline studies in both male and female α2KO mice; (2). Nicotine decreased IBAT [^18^F]FDG uptake in both males and females α2KO mice; (3). There was no sex effect between males and females either in the baseline or the nicotine treatment study. Thus, presence of α2HS receptors does not lead to a major change in the use of glucose in IBAT, but the lack of α2 receptors cause a reversal of nicotine effects in glucose metabolism in IBAT.

## Acknowledgements

Research support provided by NIH AG R01 AG 029479 (JM), AG077700 (JM), R01 DC017687 (RM), R01 AG067073 (RM) and the Undergraduate Research Opportunities Program (UROP) at University of California, Irvine. We thank Ms. Lena Qin for technical assistance.

## Author Contributions

All authors had full access to all the data in the study and are responsible for the integrity of the data and the accuracy of the data analysis. Conceptualization, J.M., RM; methodology, C.L., J.M., I.I., R.L.; validation and analysis, C.L., T.E.T., A.D.L.C., E.H.N.N., J.L.N.; investigation, J.M..; resources, I.I., R.L., R.M., J.M.; writing & editing—J.M., C.L., J.L.N., T.E.T., A.D.L.C.; supervision, J.M..; project administration, J.M..; funding acquisition, J.M. and R.M.

## Data Sharing

The data that support the findings of this study are available from the corresponding author upon reasonable request.

## References

1. Dani JA. Neuronal Nicotinic Acetylcholine Receptor Structure and Function and Response to Nicotine. International review of neurobiology. 2015 Jan 1;124:3–19. 10.1016/bs.irn.2015.07.001

2. Zhang HJ, Zammit M, Kao CM, Govind AP, Mitchell S, Holderman N, et al. Trapping of Nicotinic Acetylcholine Receptor Ligands Assayed by In Vitro Cellular Studies and In Vivo PET Imaging. The Journal of Neuroscience. 2022 Aug 26;43(1):2–13. 10.1523/JNEUROSCI.2484-21.2022

3. Whiteaker P, Wilking JA, Brown RW, Brennan RJ, Collins AC, Lindstrom JM, et al. Pharmacological and immunochemical characterization of α2* nicotinic acetylcholine receptors (nAChRs) in mouse brain. Acta Pharmacologica Sinica. 2009 Jun 1;30(6):795–804. 10.1038/aps.2009.68

4. Picciotto MR, Kenny PJ. Mechanisms of Nicotine Addiction. Cold Spring Harbor Perspectives in Medicine. 2020 Apr 27;11(5):a039610. 10.1101/csHSerspect.a039610

5. Intskirveli I, Gil S, Lazar R, Metherate R. Alpha-2 nicotinic acetylcholine receptors regulate spectral integration in auditory cortex. Frontiers in Neural Circuits. 2024 Nov 1;18. 10.3389/fncir.2024.1492452

6. Hilscher MM, Mikulovic S, Perry S, Lundberg S, Kullander K. The alpha2 nicotinic acetylcholine receptor, a subunit with unique and selective expression in inhibitory interneurons associated with principal cells. Pharmacological Research. 2023 Aug 29;196:106895. 10.1016/j.phrs.2023.106895

7. Mukherjee J, Lao PJ, Betthauser TJ, Samra GK, Pan M, Patel IH, et al. Human brain imaging of nicotinic acetylcholine α4β2* receptors using [18 F]Nifene: Selectivity, functional activity, toxicity, aging effects, gender effects, and extrathalamic pathways. Journal of Comparative Neurology. 2017 Sep 19;526(1):80–95. 10.1002/cne.24320

8. Lotfipour S, Mojica C, Nakauchi S, Lipovsek M, Silverstein S, Cushman J, et al. α2* Nicotinic acetylcholine receptors influence hippocampus-dependent learning and memory in adolescent mice. Learning & Memory. 2017 May 15;24(6):231–44. https://learnmem.cshlp.org/content/24/6/231

9. Lotfipour S, Byun JS, Leach P, Fowler CD, Murphy NP, Kenny PJ, et al. Targeted Deletion of the Mouse α2 Nicotinic Acetylcholine Receptor Subunit Gene (*Chrna*2) Potentiates Nicotine-Modulated Behaviors. The Journal of Neuroscience. 2013 May 1;33(18):7728–41. doi: 10.1523/JNEUROSCI.4731-12.2013

10. Papke RL. Functions and pharmacology of α2β2 nicotinic acetylcholine receptors; in and out of the shadow of α4β2 nicotinic acetylcholine receptors. Biochemical Pharmacology. 2024 May 10;225:116263. 10.1016/j.bcp.2024.116263

11. Liang C, Okamoto AA, Karim F, Kawauchi S, Mukherjee J. Disruption of normal brain distribution of [^18^F]nifene to a4b2* nicotinic acetylcholinergic receptors in B6129SF2/J mice and transgenic 3xTg mouse model of Alzheimer’s disease: In vivo [^18^F]Nifene PET/CT imaging studies. Neuroimage, 2025; 310: 121092. 10.1016/j.neuroimage.2025.121092

12. Liang C, Nguyen GA, Danh TB, Sandhu AK, Melkonyan LL, Syed AU, et al. Abnormal [18 F]NIFENE binding in transgenic 5xFAD mouse model of Alzheimer’s disease: In vivo PET/CT imaging studies of α4β2* nicotinic acetylcholinergic receptors and in vitro correlations with Aβ plaques. Synapse (New York, NY). 2023 Feb 23;77(3). 10.1002/syn.22265

13. Bieszczad KM, Kant R, Constantinescu CC, Pandey SK, Kawai HD, Metherate R, et al. Nicotinic acetylcholine receptors in rat forebrain that bind ^18^F-nifene: Relating PET imaging, autoradiography, and behavior. Synapse. 2012 Feb 15;66(5):418–34. 10.1002/syn.21530

14. Nguyen C, Mondoloni S, Le Borgne T, Centeno I, Come M, Jehl J, et al. Nicotine inhibits the VTA-to-amygdala dopamine pathway to promote anxiety. Neuron. 2021 Jul 8;109(16):2604–2615.e9. 10.1016/j.neuron.2021.06.013

15. Zhou M, Qiu W, Ohashi N, Sun L, Wronski ML, Kouyama-Suzuki E, et al. Deep-Learning-Based Analysis Reveals a Social Behavior Deficit in Mice Exposed Prenatally to Nicotine. Cells. 2024 Feb 1;13(3):275. DOI: 10.3390/cells13030275

16. Grant RA, Cielen N, Maes K, Heulens N, Galli GLJ, Janssens W, et al. The effects of smoking on whisker movements: A quantitative measure of exploratory behaviour in rodents. Behavioural Processes. 2016 Apr 1;128:17–23. 10.1016/j.beproc.2016.03.021

17. Michalak A, Biala G. Calcium homeostasis and protein kinase/phosphatase balance participate in nicotine-induced memory improvement in passive avoidance task in mice. Behavioural Brain Research. 2016 Sep 12;317:27–36. https://pubmed.ncbi.nlm.nih.gov/27633557/

18. Grayson RF, Henningfield JE, London ED. Intravenous nicotine reduces cerebral glucose metabolism: A preliminary study. Neuropsychopharmacology, 2003; 28: 765–772. https://www.nature.com/articles/1300106

19. Stapleton JM, Gilson SF, Wong DF, Villemagne VL, Dannals RF, Grayson RF, et al. Intravenous nicotine reduces cerebral glucose metabolism: a preliminary study. Neuropsychopharmacology. 2002 Nov 1;28(4):765–72. 10.1038/sj.npp.1300106

20. Sharma A, Brody AL. In vivo brain imaging of human exposure to nicotine and tobacco. Handbook of experimental pharmacology. 2009 Jan 1;2009(192):145–71. doi: 10.1007/978-3-540-69248-5_6

21. Yoshida T, Sakane N, Umekawa T, Kogure A, Kondo M, Kumamoto K, et al. Nicotine induces uncoupling protein 1 in white adipose tissue of obese mice. International Journal of Obesity. 1999 Jun 1;23(6):570–5. https://www.nature.com/articles/0800870

22. Baba S, Tatsumi M, Ishimori T, Lilien DL, Engles JM, Wahl RL. Effect of Nicotine and Ephedrine on the Accumulation of 18F-FDG in Brown Adipose Tissue. Journal of Nuclear Medicine. 2007 Jun 1;48(6):981–6. 10.2967/jnumed.106.039065

23. Cannon B, Nedergaard J. Brown adipose tissue: function and physiological significance. Physiological Reviews. 2004 Jan 1;84(1):277–359. 10.1152/physrev.00015.2003

24. Mirbolooki MR, Constantinescu CC, Pan ML, Mukherjee J. Quantitative assessment of brown adipose tissue metabolic activity and volume using 18F-FDG PET/CT and β3-adrenergic receptor activation. EJNMMI Research. 2011 Dec 1;1(1). https://link.springer.com/article/10.1186/2191-219X-1-30

25. Mirbolooki MR, Upadhyay SK, Constantinescu CC, Pan ML, Mukherjee J. Adrenergic pathway activation enhances brown adipose tissue metabolism: A [18F]FDG PET/CT study in mice. Nuclear Medicine and Biology. 2013 Oct 1;41(1):10–6. 10.1016/j.nucmedbio.2013.08.009

26. Somm E. Nicotinic Cholinergic Signaling in Adipose Tissue and Pancreatic Islets Biology: Revisited Function and Therapeutic Perspectives. Archivum Immunologiae et Therapiae Experimentalis. 2013 Nov 26;62(2):87–101. https://link.springer.com/article/10.1007/s00005-013-0266-6

27. Gochberg-Sarver A, Kedmi M, Gana-Weisz M, Bar-Shira A, Orr-Urtreger A. Tnfα, Cox2 and AdipoQ adipokine gene expression levels are modulated in murine adipose tissues by both nicotine and nACh receptors containing the β2 subunit. Molecular Genetics and Metabolism. 2012 Aug 17;107(3):561–70. 10.1016/j.ymgme.2012.08.012

28. Jun H, Yu H, Gong J, Jiang J, Qiao X, Perkey E, et al. An immune-beige adipocyte communication via nicotinic acetylcholine receptor signaling. Nature Medicine. 2018 May 21;24(6):814–22. 10.1038/s41591-018-0032-8

29. Kim J. Association of CHRNA2 polymorphisms with overweight/obesity and clinical characteristics in a Korean population. Clinical Chemistry and Laboratory Medicine. 2008 Jan 1;46(8). 10.1515/CCLM.2008.230

30. Easwaramoorthy, B., Pichika, R., Collins, D., Potkin, S.G., Leslie, F.M. and Mukherjee, J.: Effect of acetylcholinesterase inhibitors on nicotinic a4b2 receptor PET radiotracer, ^18^F-Nifene: A measure of acetylcholine competition. Synapse, 61: 29–36, 2007. 10.1002/syn.20338

31. Coleman RA, Liang C, Patel R, Ali S, Mukherjee J. Brain and Brown Adipose Tissue Metabolism in Transgenic Tg2576 Mice Models of Alzheimer Disease Assessed Using 18F-FDG PET Imaging. Molecular imaging. 2017 Jan 1;16:1–9. 10.1177/1536012117704557

32. Mondal R, Campoy ADT, Liang C, Mukherjee J. 18 F]FDG PET/CT Studies in Transgenic Hualpha-Syn (A53T) Parkinson’s Disease Mouse Model of α-Synucleinopathy. Frontiers in Neuroscience. 2021 Jun 15;15(Suppl 15). 10.3389/fnins.2021.676257

33. Mineur YS, Ernstsen C, Islam A, Lefoli Maibom K, Picciotto MR. Hippocampal knockdown of α2 nicotinic or M1 muscarinic acetylcholine receptors in C57BL/6J male mice impairs cued fear conditioning. Genes, Brain and Behavior. 2020 Jul 1;19(6). 10.1111/gbb.12677

34. Mosconi L, Berti V, Glodzik L, Pupi A, De Santi S, De Leon MJ. Pre-Clinical Detection of Alzheimer’s Disease Using FDG-PET, with or without Amyloid Imaging. Journal of Alzheimer’s Disease. 2010 May 26;20(3):843–54. 10.3233/JAD-2010-091504

35. Andersen JV, Mckenna MC. Neurons in Need: Glucose, but Not Lactate, Is Required to Support Energy-Demanding Synaptic Transmission. Journal of neurochemistry. 2025 Oct 1;169(10). 10.1111/jnc.70255

36. Bouter C, Irwin C, Franke TN, Beindorff N, Bouter Y. Quantitative Brain Positron Emission Tomography in Female 5XFAD Alzheimer Mice: Pathological Features and Sex-Specific Alterations. Frontiers in Medicine. 2021 Nov 26;8(Suppl 2). 10.3389/fmed.2021.745064

37. Gandhi A, Tang R, Seo Y, Bhargava A. Organ-Specific Glucose Uptake: Does Sex Matter? Cells. 2022 Jul 16;11(14):2217. 10.3390/cells11142217

38. Feng B, Cao J, Yu Y, Yang H, Jiang Y, Liu Y, et al. Gender-Related Differences in Regional Cerebral Glucose Metabolism in Normal Aging Brain. Frontiers in Aging Neuroscience. 2022 Feb 10;14. 10.3389/fnagi.2022.809767

39. Hu Y, Xu Q, Li K, Zhu H, Qi R, Zhang Z, et al. Gender Differences of Brain Glucose Metabolic Networks Revealed by FDG-PET: Evidence from a Large Cohort of 400 Young Adults. PLoS ONE. 2013 Dec 17;8(12):e83821. 10.1371/journal.pone.0083821

40. Yoshizawa H, Gazes Y, Stern Y, Miyata Y, Uchiyama S. Characterizing the normative profile of 18F-FDG PET brain imaging: Sex difference, aging effect, and cognitive reserve. Psychiatry Research: Neuroimaging. 2013 Oct 31;221(1):78–85. 10.1016/j.pscychresns.2013.10.009

41. Vargas-Medrano J, Carcoba LM, Vidal Martinez G, Mulla ZD, Diaz V, Ruiz-Velasco A, et al. Sex and diet-dependent gene alterations in human and rat brains with a history of nicotine exposure. Frontiers in psychiatry. 2023 Feb 10;14. 10.3389/fpsyt.2023.1104563

42. Torigian DA, Green-Mckenzie J, Liu X, Shofer FS, Werner T, Smith CE, et al. A Study of the Feasibility of FDG-PET/CT to Systematically Detect and Quantify Differential Metabolic Effects of Chronic Tobacco Use in Organs of the Whole Body—A Prospective Pilot Study. Academic Radiology. 2016 Oct 18;24(8):930–40. 10.1016/j.acra.2016.09.003

43. Tong LQ, Sui YF, Jiang SN, Yin YH. The Association Between Lung Fluorodeoxyglucose Metabolism and Smoking History in 347 Healthy Adults. Journal of Asthma and Allergy. 2021 Mar 31;14(Suppl 17):301–8. 10.2147/JAA.S302602

44. Martínez-Martos JM, Cantón-Habas V, Rich-Ruíz M, Reyes-Medina MJ, Ramírez-Expósito MJ, Carrera-González MDP. Sexual and Metabolic Differences in Hippocampal Evolution: Alzheimer’s Disease Implications. Life (Basel, Switzerland). 2024 Nov 26;14(12):1547. 10.3390/life14121547

45. Van Sinderen ML, Steinberg GR, Jørgensen SB, Honeyman J, Chow JD, Herridge KA, et al. Effects of Estrogens on Adipokines and Glucose Homeostasis in Female Aromatase Knockout Mice. PLoS ONE. 2015 Aug 28;10(8):e0136143. 10.1371/journal.pone.0136143

46. Biegon A, Kim SW, Logan J, Hooker JM, Muench L, Fowler JS. Nicotine Blocks Brain Estrogen Synthase (Aromatase): In Vivo Positron Emission Tomography Studies in Female Baboons. Biological Psychiatry. 2010 Feb 25;67(8):774–7. 10.1016/j.biopsych.2010.01.004

47. Biegon A, Alia-Klein N, Fowler JS. Potential Contribution of Aromatase Inhibition to the Effects of Nicotine and Related Compounds on the Brain. Frontiers in Pharmacology. 2012 Nov 6;3(Suppl 1). 10.3389/fphar.2012.00185

48. Dubol M, Immenschuh J, Jonasson M, Takahashi K, Niwa T, Hosoya T, et al. Acute nicotine exposure blocks aromatase in the limbic brain of healthy women: A [11C]cetrozole PET study. Comprehensive Psychiatry. 2023 Mar 5;123:152381. 10.1016/j.comppsych.2023.152381

49. Carpentier AC, Blondin DP, Haman F, Richard D. Brown Adipose Tissue-A Translational Perspective. Endocrine reviews. 2022 May 29;44(2):143–92. 10.1210/endrev/bnac027

50. Trujillo ME, Scherer PE. Adipose Tissue-Derived Factors: Impact on Health and Disease. Endocrine Reviews. 2006 Oct 20;27(7):762–78. 10.1210/er.2006-0033

51. Seoane-Collazo P, Liñares-Pose L, Rial-Pensado E, Romero-Picó A, Moreno-Navarrete JM, Martínez-Sánchez N, et al. Central nicotine induces browning through hypothalamic \u03ba opioid receptor. Nature Communications. 2019 Sep 6;10(1). 10.1038/s41467-019-12004-z

52. Qin R, Zhang Y, Xu S, Mei Y, Jin G, Mi Y, et al. Effects of Nicotine Doses and Administration Frequencies on Mouse Body Weight and Adipose Tissues. Nicotine & tobacco research : official journal of the Society for Research on Nicotine and Tobacco. 2024 Sep 5;27(3). 10.1093/ntr/ntae208

53. De Paula Faria D, Da Silva Vera CC, Marques FLN, Sapienza MT. Repeatability of brown adipose tissue activation measured by [18F]FDG PET after beta3-adrenergic stimuli in a mouse model. Nuclear medicine and biology. 2023 Sep 29;126–127:108390. 10.1016/j.nucmedbio.2023.108390

54. Mukherjee J, Baranwal A, Schade KN. Classification of Therapeutic and Experimental Drugs for Brown Adipose Tissue Activation: Potential Treatment Strategies for Diabetes and Obesity. Current diabetes reviews. 2016 Oct 26;12(4):414–28. https://pmc.ncbi.nlm.nih.gov/articles/PMC5425649/

55. Li Y, Mao J, Chai G, Zheng R, Liu X, Xie J. Neurobiological mechanisms of nicotine’s effects on feeding and body weight. Neuroscience and Biobehavioral Reviews. 2025;169:106021. 10.1016/j.neubiorev.2025.106021

56. Kooijman S, van den Heuvel JK, Rensen PCN. Neuronal control of brown fat activity. Cell Press. 2015; 26(11):657. 10.1016/j.tem.2015.09.008

